# Molecular mechanism of SARS-CoV-2 inactivation by temperature

**DOI:** 10.1101/2020.10.16.343459

**Authors:** Didac Martí, Juan Torras, Oscar Betran, Pau Turon, Carlos Alemán

## Abstract

Recent studies have shown that SARS-CoV-2 virus can be inactivated by effect of heat, even though, little is known about the molecular changes induced by the temperature. Here, we unravel the basics of such inactivation mechanism over the SARS-CoV-2 spike glycoprotein by executing atomistic molecular dynamics simulations. Both the *closed down* and *open up* states, which determine the accessibility to the receptor binding domain, were considered. Results suggest that the spike undergoes drastic changes in the topology of the hydrogen bond network while salt bridges are mainly preserved. Reorganization in the hydrogen bonds structure produces conformational variations in the receptor binding subunit and explain the thermal inactivation of the virus. Conversely, the macrostructure of the spike is preserved at high temperature because of the retained salt bridges. The proposed mechanism has important implications for engineering new approaches to inactivate the SARS-CoV-2 virus.

## Introduction

The inactivation of SARS-CoV-2 has become a major objective as the spread of the beta coronavirus (β-CoV) has triggered a global health crisis with high social (more than 37 million infected people and more than one million deaths worldwide as of September 29^th^) and economic (global GDP contraction up to 5.2 % for the year 2020) impacts (World Health Organization; World Bank). Virus inactivation can be achieved using different strategies based on chemical, biological and physical treatments. Biocide chemical agents and surfactants are frequently used to disinfect inanimate surfaces (Kampf *et al.* 2020a; Smith et al. 2020). Furthermore, virus in culture media are deactivated by chemicals as TRIzol^®^, Formalin (formaldehyde) and β-propiolactone (Jureka *et al.* 2020). Biological treatments, in the format of vaccines, are the most effective to fight the virus in living organisms, even though their successful development is sometimes limited by a combination of economic factors, regulatory environment and the empirical nature of modern vaccine discovery (Loomis and Johnson, 2015; Rappuoli *et al.* 2014; Tannock *et al.* 2020). Among physical treatments, cold plasma (Filipić *et al.*, 2020), far-UVC light (Buonanno *et al.*, 2020) and membrane filtration (Shirasaki *et al.*, 2017) have been described to eliminate virus in surfaces, air and water, respectively, even though thermal inactivation is the most extensively used treatment when possible (Abraham *et al.*, 2020; Cimolai, 2020; Hu *et al.*, 2020; Jureka *et al.* 2020; Kampf *et al.*, 2020b; Pastorino *et al.* 2020; Yap *et al.*, 2020).

Focusing on the inactivation by temperature, it has been reported that SARS-CoV-2 was highly stable at low temperatures (*i.e.* 4 °C up to 14 days; Chin *et al.*, 2020) but it is sensitive to heat. Indeed, several minimal inactivation temperatures, which are comprised between 56 °C (45 min for total inactivation) and 100 °C (< 5 min for total inactivation; Jureka *et al.* 2020), have been reported (Abraham *et al.*, 2020; Cimolai, 2020; Hu *et al.*, 2020; Jureka *et al.* 2020; Kampf *et al.*, 2020b; Pastorino *et al.* 2020; Yap *et al.*, 2020). On the other hand, the infectivity of SARS-CoV-2 in solution was strongly reduced (up to 100-fold), even though the virus remained similarly infective in surfaces at 4 and 30 °C (Kratzel *et al.*, 2020).

Temperature is known to promote changes in the molecular structure of biomacromolecules (*i.e.* nucleic acids, proteins, lipids) until affecting their functionality. As part of natural evolution, such alterations have been used by microorganisms as an adaptation mechanism to environmental changes, for instance by variations in thermal labile hydrogen bonds that connect the strands of nucleic acids and proteins (Sengupta and Garrity, 2013). In the particular case of proteins, temperature is known to induce variations of the secondary and tertiary structure, causing structural alterations that modify their stability and their role in regulating cellular processes, signal transduction and intrinsic enzymatic properties. However, the response of proteins to changes in the temperature conditions can be very different. For example, some proteins have high thermal stability while others can unfold or even denature at moderate temperatures (Dong *et al.*, 2018; Julió Plana *et al.*, 2019; Lopes-Rodrigues *et al.*, 2018).

In this work we aim to unravel the effect of the temperature on the molecular structure of the SARS-CoV-2 spike glycoprotein, which is a homotrimer with three monomers with identical primary structure, named chain A, B and C. Coronaviruses use the spike to bind cellular receptors, triggering a cascade of events that leads to cell entry (Song *et al.*, 2018; Zhou *et al.*, 2020). The spike protein is reported to bind the cellular receptor human angiotensin-converting enzyme 2 (ACE2) that mediates the fusion of the viral and cellular membrane and facilitates the virus introduction inside the cell (Ou *et al.*, 2020). Each spike monomer (180 kDa) contains 1273 amino acids (aa) and consists of a signal peptide (aa 1-13) and two subunits, named S1 (aa 14-685) and S2 (aa 686-1273), which are responsible for receptor binding and membrane fusion, respectively. The S1 subunit involves the N-terminal domain (NTD; aa 14-305) and the receptor binding domain (RBD; 319-541). The latter consists of a core region with 5 β-pleated sheets (β1, β2, β3, β4 and β7) organized in antiparallel model and the receptor-binding motif with two short β-pleated sheets (β5 and β6), loops and alpha helices (α4 and α5; Shang *et al.*, 2020). A total of three cysteine residue pairs provide stability to the core and an additional cysteine residue pair helps in connecting the distal end of the receptor-binding motif (Lan *et al.*, 2020). The S2 subunit has a key role in the membrane fusion (Kirchdoerfer *et al.*, 2016). Thus, after the initial interaction of the RBD of the S1 subunit and the peptidase domain of ACE2, the fusion with the host cell membrane continues through the interaction between the heptad repeat 1 domain (HR1; aa 912-984), the central helix (CH; aa 985-1035), the connector domain (CD; aa 1076-1141), the heptad repeat 2 domain (HR2; aa 1163-1213) of S2 to form a helix bundle fusion core (Jan Bosch *et al.*, 2004), which is extra stabilized by a short residue sequence (named fusion peptide of S2 or FP; aa 788-806; Xia *et al.*, 2020). Furthermore, another important structural aspect of the SARS-CoV-2 spike is the identification of two states for the hinge-like conformation of the RBD of S1 from chain B (Walls *et al.*, 2020; Wrapp *et al.* 2020). These are: 1) the *closed down* state (Figure 1a) in which the RBD of S1 covers the apical region of S2 near the C-terminus of HR1 (*i.e.* the receptor binding for interacting with ACE2 is buried); and 2) the *open up* (Figure 1b) state in which the RBD is dissociated from the central axis of S2 and the NTD of S1 (*i.e.* the receptor binding motifs are exposed).

**Figure 1.**
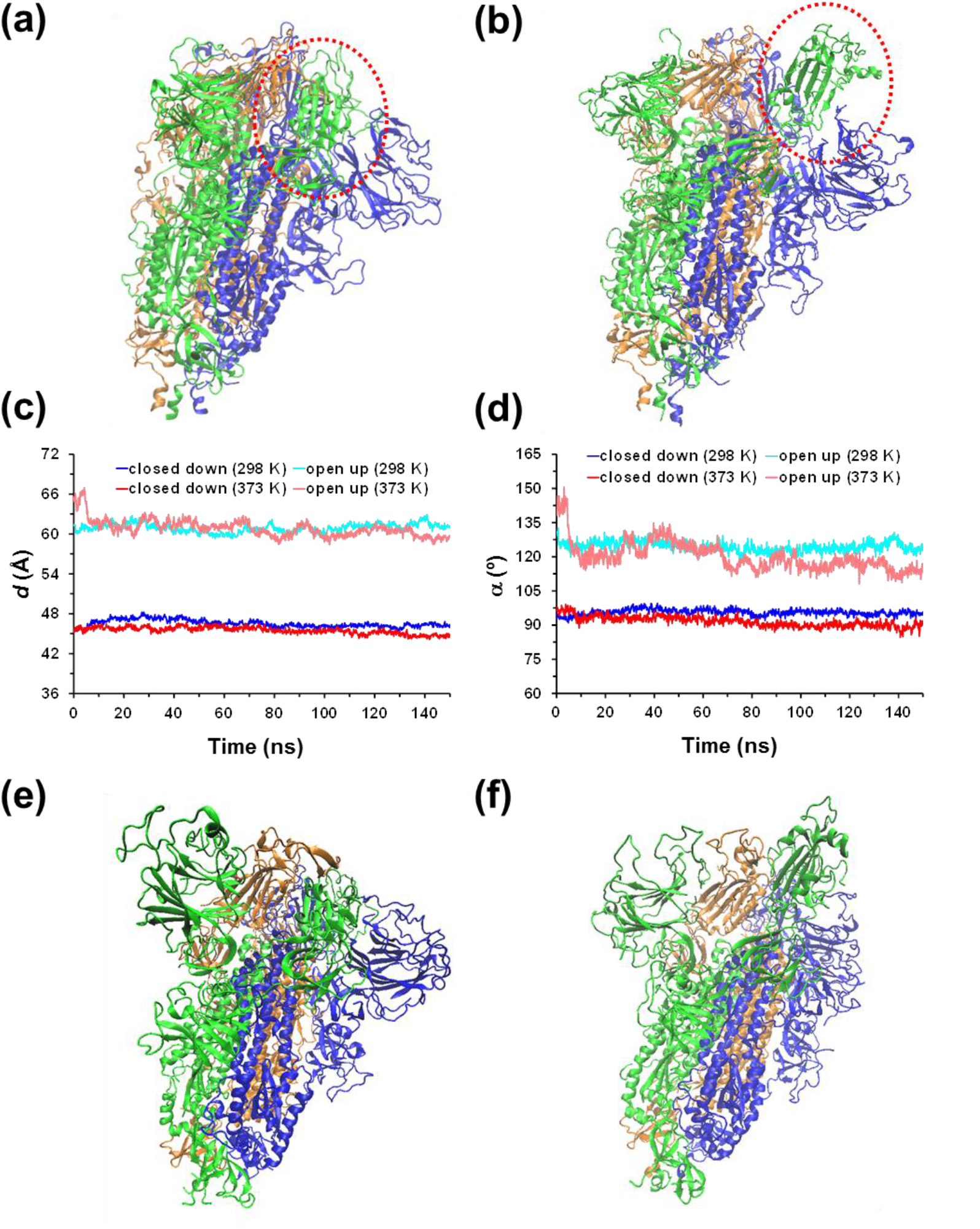
(a) Closed down and (b) open up conformational states of the SARS-CoV-2 homotrimeric spike protein. The main difference between such two states is marked by the red dashed circle. For the MD simulations conducted at 298 and 373 K, temporal evolution of the geometric parameters used to identify the conformational states of the protein: (c) distance (*d*) between the centre of mass of the RBD from chain B to the centre of mass of three monomers; and (d) hinge angle (*α*) formed by the center of mass of the RBD of chain B, the center of mass of the rest of chain B, and the center of mass of the first residue after the RBD. For the (e) closed down and (f) open up states, structure of the spike protein at the end of the MD simulation at 373 K.

This study is focused on exploring the molecular structure of the spike glycoprotein of SARS-CoV-2 at different temperatures comprised within 25 °C and 100 °C using atomistic computer simulations. More specifically, atomistic Molecular Dynamics (MD) simulations have been conducted on the homotrimeric protein in aqueous solution considering both the closed down and the open up conformational states. MD is a computational tool that captures time-dependent conformational changes at various conditions by calculating inter-atomic forces through solvent Newton’s second law. Moreover, this *in silico* methodology has been used to trace the atomic level contacts and interactions, which cannot be captured experimentally, and unveil a molecular mechanism for the virus inactivation. Such mechanism is intended to contribute to the development of new inactivation strategies by means of physical or chemical treatments specifically designed to disable such key interactions identified through the functional sites of the virus.

## Results

The atomic coordinates of the homotrimeric spike glycoprotein of SARS-CoV-2 in the closed down and open up conformational states were taken from the Protein Data Bank (PDB; entry 6vxx and 6vyb, respectively, from cryo-electron microscopy structures of the SARSCoV-2 spike ectodomain trimer; Walls *et al.*, 2020). Missing protein segments were built using the UCSF Chimera program (Pettersen *et al.*, 2004), which used the Modeller algorithm (Webb and Sali, 2016) to fill with the FASTA amino acids sequence while maintaining the crystallographic structure fixed. The different models generated for each conformational state were assessed using Z-DOPE (Discrete Optimized Protein Energy), a normalized atomic distance dependent statistical potential based on known protein structures. For each conformational state, the model scored with the lowest Z-DOPE, which was lower than –1 for both cases, was selected for MD simulations. After adding the hydrogen atoms and forming the disulphide bonds, the structures were solvated, thermalized and equilibrated using the protocol described in the Methods section (Supplementary Information). Finally, 150 ns long MD production runs in an NVT ensemble were conducted at 298, 310, 324, 338, 358 and 373 K using the Amber18 program (Case *et al.*, 2018). In order to ensure the repeatability of the observed tendencies, replica simulations were performed at defined temperatures by changing the initial velocities. Accordingly, a total of 3.6 μs (150 ns × 2 conformational states × 6 temperatures × 2 replicas) were simulated for such a large system, which involved more than half million atoms.

The influence of the temperature on the closed down and open up states is quantitatively analyzed in Figure 1c-d, which compares the temporal evolution of two different parameters calculated using data derived from simulations at 298 K and 373 K. These parameters, which have been defined to clearly differentiate between the two conformational states as well as to be able to identify the presence of intermediate states, are: 1) the distance (*d*) between the centre of mass of the RBD of chain B to the centre of mass of the three monomers; and 2) the hinge angle (*α*) formed by the center of mass of the RBD from chain B, the center of mass of the rest of chain B (*i.e.* excluding the RBD), and the center of mass of the first residue after the RBD. As it was expected, both parameters remain practically unaltered (less than 1.5% and 5.2% for *d* and *α*, respectively) along the MD trajectories at 298 K, the average values being *d* = 46.5 ± 0.5 Å and *α* = 95.6 ± 1.4° for the closed down and *d* = 60.9 ± 0.6 Å and *α* = 124.4 ± 2.2° for the open up. Interestingly, the average values obtained at 373 K, *d* = 45.5 ± 0.5 Å and *α* = 91.6 ± 2.3° for the closed down and *d* = 60.8 ± 1.3 Å and *α* = 120.2 ± 6.3° for the open up, are very similar compared to the previous one. Although the fluctuations around the average values are slightly higher at 373 K, especially for the open up conformation, results indicate the two states are unambiguously maintained during the whole simulations. Accordingly, the thermal energy gained at 373 K is not enough to destabilize the open up state and initiate a transition process towards the closed down. The same behavior was obtained for the rest of the studied temperatures (not shown).

Although detailed energetic analyses of the closed down and open up conformations are out of the scope of this work, we observed that the former state is stabilized with respect to the latter one, independently of the temperature. This stability order is in agreement with experimental observations (Pallesen *et al.* 2017; Walls *et al.*, 2020; Yuan *et al.*, 2017). Even though the two conformational states are maintained throughout all the trajectories, temperature affects the spike structure, causing changes that, after a certain threshold, cause the inactivation of the virus (Abraham *et al.*, 2020; Cimolai, 2020; Hu *et al.*, 2020; Jureka *et al.* 2020; Kampf *et al.*, 2020b; Pastorino *et al.* 2020; Yap *et al.*, 2020). These changes are illustrated in Figure 1e-f, which show the closed down and open up conformations obtained at the end of the MD trajectories at 373 K. As can be seen, the structures present significant differences to naked eye with respect to those shown in Figure 1a-b, which were used as a starting point. These differences affect not only the secondary structure, but also the quaternary structure, which describes the interchain assembly between the three monomers. Thus, visual analysis of the open up conformation suggests that one of the monomers undergoes densification in certain regions due to structural deformations occurring at 373 K (blue monomer in Figure 1f).

The influence of the temperature on the interchain assembly was analyzed by exploring the temporal evolution of the interchain distances (*i.e.* distance between two monomers), which was calculated as the distance between mass centres. Figure 2a-b compares the A-B, B-C and A-C interchain distances at 298 and 373 K for the closed down and open up states. For the closed down state, the three interchain distances remain within the 32-34 Å interval, independently of the temperature. Instead, the open up state shows different interchain distances, even at room temperature. For example, at 298 K the values averaged over the last 75 ns of simulation are: d(A-B) = 37.1 ± 0.2 Å, d(B-C) = 35.6 ± 0.2 Å and d(A-C) = 31.9 ± 0.2 Å, reflecting that monomer A is closer to C than to B by ~9%. At 373 K, d(A-B) and d(B-C) converge to practically the same value (*i.e.* 35.4 ± 0.4 Å and 35.2 ± 0.3 Å, respectively), whereas d(A-C) decreases to 29.6 ± 0.2 Å. The latter represents a reduction of 16% with respect to the other two values and a reduction of 7% with respect to the distance at 298 K.

**Figure 2.**
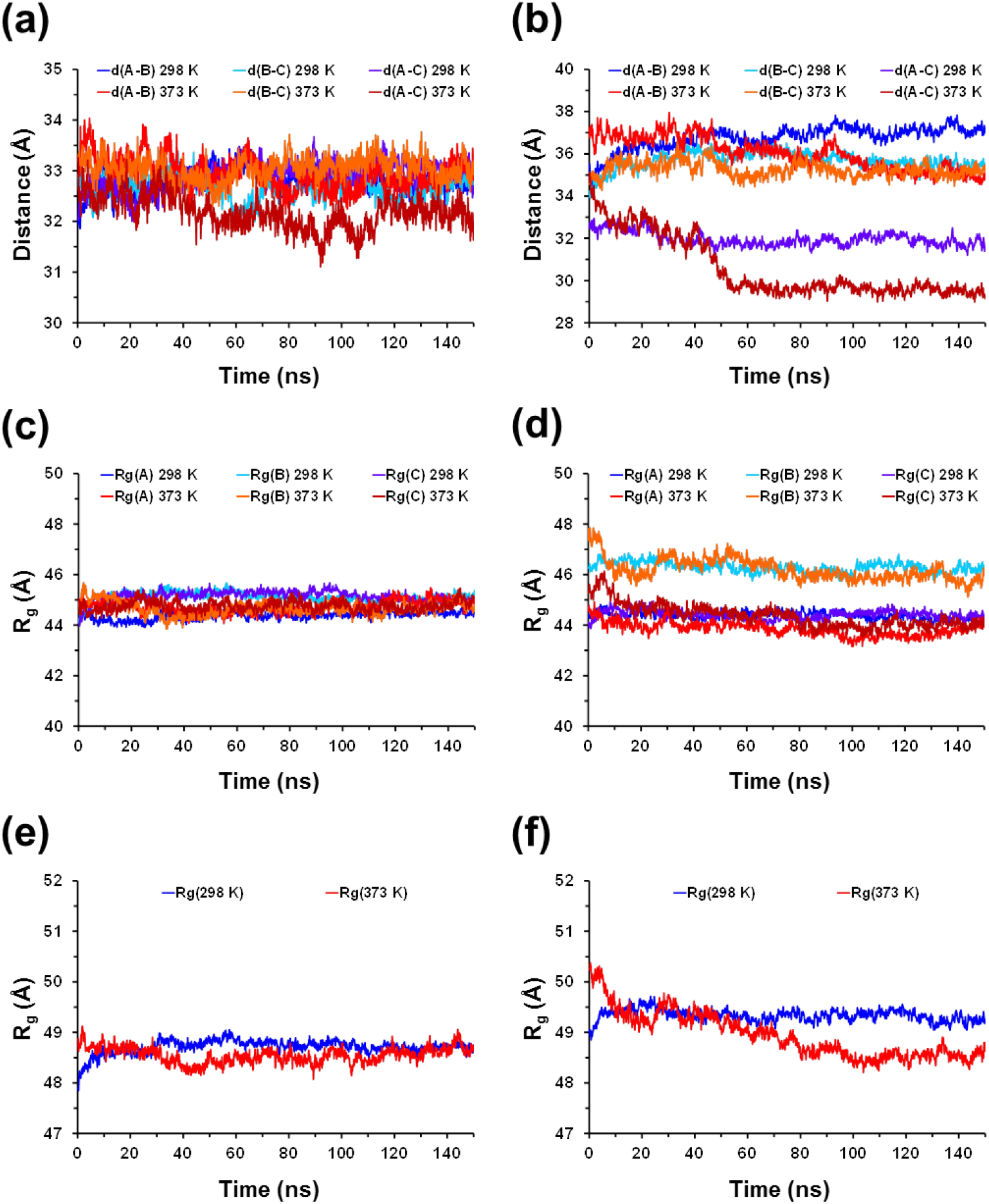
For the (a, c, e) closed down and (b, d, f) open up states, temporal evolution at 298 and 373 K of: (a, b) the inter-chains distances calculated with respect to the centre of mass of each chain; (c, d) the radius of gyration of each chain; and (e, f) the radius of gyration of the whole supramolecular assembly formed by the three assembled chains.

In order to ascertain if the effect on the assembly of the chains is associated to a change in the global shape of the individual monomers and of the whole homotrimeric construct, the dynamics of the radius of gyrations (R_g_) was followed. The R_g_ for the individual chains does not exhibit abrupt changes when the temperature increases from 298 to 373 K (Figure 2c-d). For the closed down state, the R_g_ is ~45 Å at the two analyzed temperatures. The R_g_ of chains A and C is ~44 Å for the open up state and slightly larger for chain B (~46 Å), even though the influence of the temperature in such values is very small (*e.g.* the R_g_ of monomer B averaged over the last 75 ns of the trajectory at 298 and 373 K is 46.2 ± 0.2 and 45.9 ± 0.2 Å, respectively). Similarly, although the effect of raising the temperature from 298 K to 373 K on the R_g_ of the whole assembly is slightly more pronounced for the open state (*i.e.* from 49.3 ± 0.1 Å to 48.6 ± 0.1 Å) than for the closed one (*i.e.* from 48.7 ± 0.1 Å to 48.5 ± 0.1 Å), changes do not reflect an important variation in shape of the supramolecular assembly (Figure 2e-f).

All atom root mean square deviation (RMSD) and root mean square fluctuation (RMSF) plots determined from the MD trajectories for the closed down and open up conformational states at all the examined temperatures are illustrated in Figures 3a-b and 3c-d, respectively. At the beginning of each trajectory, we observe a rapid increase in the RMSD, independently of the temperature and conformational state. However, the structure of the spike reaches a steady state after a few tenths of ns, even at the higher temperatures, corroborating that the simulated states are well equilibrated. The RMSDs recorded at 298, 310, 324 and 338 K are relative similar, around 4-5 Å, indicating moderate structural rearrangements, whereas the maximum values obtained at 358 and, especially, 373 K are consistent with more severe structural changes. Specifically, the RMSD values averaged over the last 50 ns of the simulations at 358 and 373 K are 6.8 and 8.3 Å, respectively, for the close down state and 8.8 and 12.7 Å, respectively, for the open up state.

**Figure 3.**
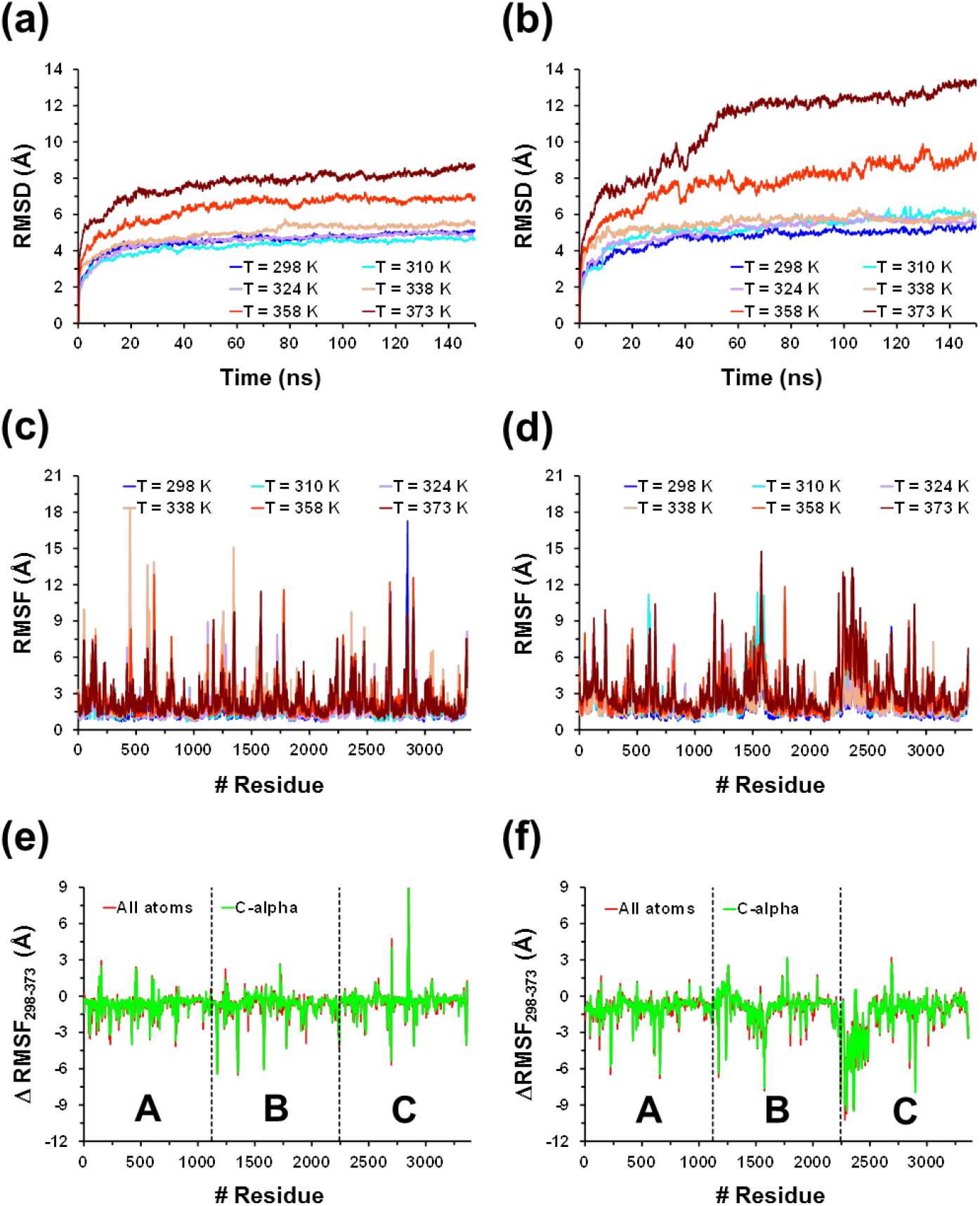
For the (a, c, e) closed down and (b, d, f) open up states of SARS-CoV-2 homotrimeric spike protein: (a, b) Temporal evolution of the root mean square deviation (RMSD) and (c, d) root mean square fluctuation (RMSF) analyses calculated using all atoms for the MD simulations conducted at 298, 310, 324, 338, 358 and 373 K; and (e, f) difference between the RMSFs obtained at 298 and 373 K (ΔRMSF= RMSF(298 K) – RMSF(373 K) calculated using all the atoms (red profile) and the C^α^ atom (green profile) of each residue. The black dashed lines separate the three monomers (A, B and C) of the spike protein.

In order to compare the overall flexibility of the two conformational states at the different temperatures, the RMSF was computed considering all atoms of each residue from the trimeric protein (Figure 3c-d). As it was expected, fluctuations increase with temperature for the two conformational states. However, increments were relatively small for temperatures ≤ 338 K (*i.e.* average value grew from 1.7 / 1.9 Å (298 K) to 2.2 / 2.3 Å (338 K) for the closed down / open up states), increasing to values ~3 Å in average at 358 and 373 K. This feature is clearly illustrated in Figure 3e-f, which represents the difference between the RMSFs calculated at 298 and 373 K considering all the atoms of each residue (ΔRMSF_298-373_).

Figure 3e-f represents the difference between the RMSFs calculated at 298 and 373 K considering both all the atoms and only the C^α^ atom of each residue (ΔRMSF_298-373_). The profiles calculated using all atoms and the C^α^ atom are very similar, independently of the conformational state, indicating that the increment of temperature has a deep impact on the backbone organization and, therefore, on the secondary structure. Moreover, the effect is apparently different for the three chains of the homotrimer (split by dashed black lines in Figure 3e-f). Thus, the thermal energy generated by heating from 298 to 373 K produces more fluctuations in the C chain for both the closed down and the open up states.

In order to look in detail the effect of the temperature on the different domains for the two states, Figure 4 compares the RMSF calculated at 298 and 373 K considering all atoms of each residue of chains A, B and C. As is shown, the position of the following domains is indicated for each subunit of each chain: 1) the NTD and the RBD from the S1 subunit; and 2) the FD, the HR1, the CH and the CD from the S2 subunit. For chain A, the S1 subunit is much more affected by the temperature than S2, reflecting that β-sheet rich domains, such as the NTD and the RBD, are more flexible than the helical rich domains located at S2. The flexibility of S1 is higher for the open up than for the closed down, as is evidenced by the fact that the RMSFs values reached at 373 K are, in general, higher for former state than for the latter one. Instead, the rigidity of S2 is similar for the two states with exception of HR1 domain, which shows large fluctuations at the central region for the closed down state. Inspection of the RMSFs achieved at intermediate temperatures indicates that the fluctuations reached by the S1 subunit of chain A increases progressively with heating. This is illustrated in Figure S1, which displays the RMSF calculated at 324 and 358 K

**Figure 4.**
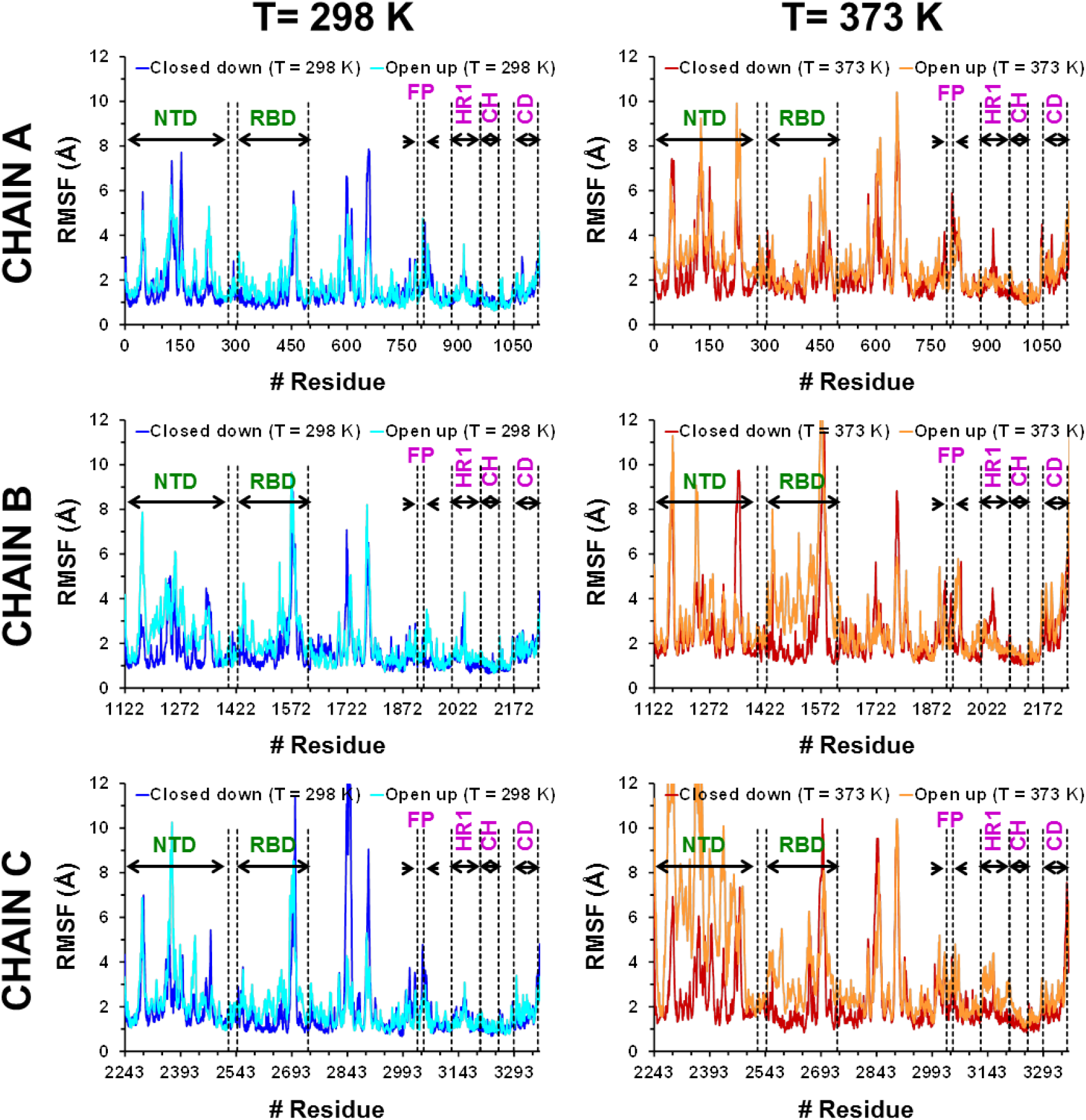
For the three chains of the homotrimeric spike protein in the closed down and the open up states: root mean square fluctuation (RMSF) analyses calculated using all atoms for the molecular dynamics simulations at 298 K (left) and 373 K (right). The following domains are indicated in each graphic: the N-terminal and the receptor binding domain (NTD and RBD, respectively) from S1 subunit (marked in green), and the fusion peptide (FD), the heptapeptide repeat sequence 1 (HR1), the central helix (CH), the connector domain (CD) from S2 subunit (marked in violet).

The behavior shown by chain B is similar to that of chain A (Figures 4 and S1), the S1 subunit being much more influenced by the increase in temperature than S2. However, it should be noted that the fluctuations observed at 373 K in the NTD and RBD domains are much higher for the B chain than for the A one, indicating that in this case the flexibility of the S1 subunit is greater. This increase in the flexibility of S1 is particularly striking for the open state that reaches RMSF values close to 12 Å (NTD) or even higher (RBD). The effect of temperature in the open state is even more pronounced for the C chain, which shows a marked destabilization of the NTD at 373 K. It is worth noting that the thermal disruption of the NTD for the open up conformation also occurs at 358 K (Figure S1), while the RSMF profiles of the two states at 324 K are similar to that calculated at 298 K.

In order to assess how the temperate affects the network of hydrogen bonds, a detailed analysis on this specific interaction was conducted. The geometric parameters and the cut-off values used to define the D–H···A interaction (where A is an acceptor atom, D a donor heavy atom, and H a hydrogen atom) as hydrogen bond are both the ∠DHA angle, which must be greater than 135°, and the D···A distance, that must be lower than 3.0 Å. Figure S2a compares the temporal evolution of the total amount of hydrogen bonds existing and reorganized in the homotrimeric protein for the closed down and open up states at both 298 and 373 K. The profiles obtained at both temperatures are very similar, the average values for the closed down / open up conformations are 1566 ± 26 / 1535 ± 26 and 1538 ± 30 / 1557 ± 27 at 298 and 373 K, respectively. Moreover, intermediate temperatures exhibit a similar behavior (*e.g.* the amount of hydrogen bonds at 338 K is 1570 ± 28 and 1560 ± 27 for the closed down and open up, respectively). Detailed analyses of the hydrogen bonds involving residues from the RBD (Figure S2b) and NTB (Figure S2c) of chains A, B and C, which are the domains that exhibit the higher fluctuations at 373 K (Figure 4), confirm that the change in terms of the number of hydrogen bonds is not significant when the temperature increases from 298 to 373 K. For example, for the open up state the average number of hydrogen bonds in the RBD of A, B and C chains is 72 ± 6 / 72 ± 6, 78 ± 7 / 80 ± 8 and 76 ± 6 / 74 ± 6 at 298 K / 373 K, respectively.

Figure 5a shows the hydrogen bonding topology map at the end of the simulation at 373 K for both the closed down and open up states. Although a large number of hydrogen bonds are detected for the two states, only a few ones involve the same residues of those existing at beginning of the simulations (marked by red dots). Thus, a large number of newly formed hydrogen bonds implies a significant reorganization of the topological map (*i.e.* at the end of the simulation, the residue that involves the donor or acceptor atom in the hydrogen bond is different from the one at the beginning). Conversely, original hydrogen bonds are mostly preserved during the dynamics run at 298 K, as is shown in Figure 5b for both studied conformational states. The hydrogen bond topology through the 358 K trajectory remains roughly the same as 298 K.. The changes induced by temperature in the RBD region of the open up state are illustrated in Figures 5c and S3, which compare the structure of such region in the initial and the last snapshot after 150 ns at 373 K and 298 K, respectively. As is shown, structural re-arrangements are very significant at the highest temperature. Similar features are shown in Figure S4 for the closed down state.

**Figure 5.**
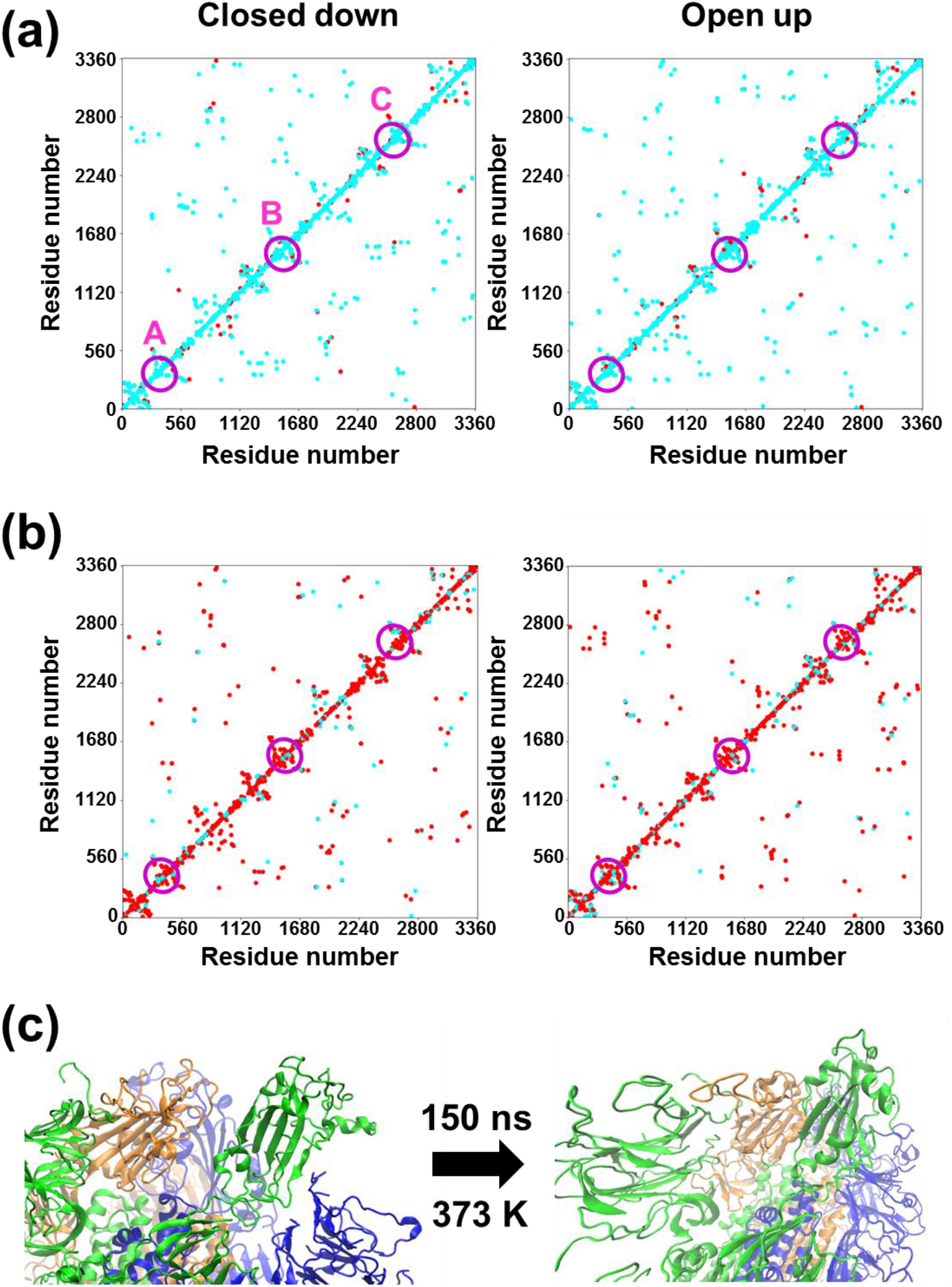
Topology maps for the hydrogen bonds found at the end of the simulations at (a) 373 K and (b) 298 K for the closed down and open up states. The *x*-and *y*-axes represent the residue number containing the hydrogen bonding donor and acceptor atoms, respectively. Red dots indicate hydrogen bonds present at the starting structure that are maintained at the end of the simulation (*i.e.* the two residues involved in the hydrogen bond do not change), while light blue dots correspond to the hydrogen bonds that were not present at the starting conformation (*i.e.* the residue involving the hydrogen bonding donor and/or acceptor atoms change). The empty purple circles indicate the position of the RBD in the three chains of the homotrimer. (c) Representation of the region containing the RBD domains at the beginning and the end of the simulation for the open up state at 373 K.

The formation and temporal evolution of unspecific salt bridges, defined by an ion pair distance ≤ 4 Å, have been investigated considering the large amount of charged residues existing in the primary structure of the modelled protein (*i.e.* 49 Lys, 37 Arg, 15 His, 40 Glu and 53 Asp per monomer). At 298 K the average amount of salt bridges is 178 ± 9 and 160 ± 7 for the closed down and open up states, respectively, increasing to 198 ± 10 and 188 ± 9, respectively, at 373 K (Figure S5). Inspection of the topology maps for the salt bridges at the end of the simulations, which are displayed in Figure 6, indicates that 268 (closed down) / 246 (open up) and 254 (closed down) / 224 (open up) of the initial interactions are retained after 150 ns at 298 and 373 K, respectively. Thus, around 73% / 66% and 69% / 61 % of the initial salt bridges are preserved at 298 and 373 K, respectively. This fact suggests that such strong interactions, which are mainly of electrostatic nature, are responsible of the approximate conservation of the general shape of the spike protein.

**Figure 6.**
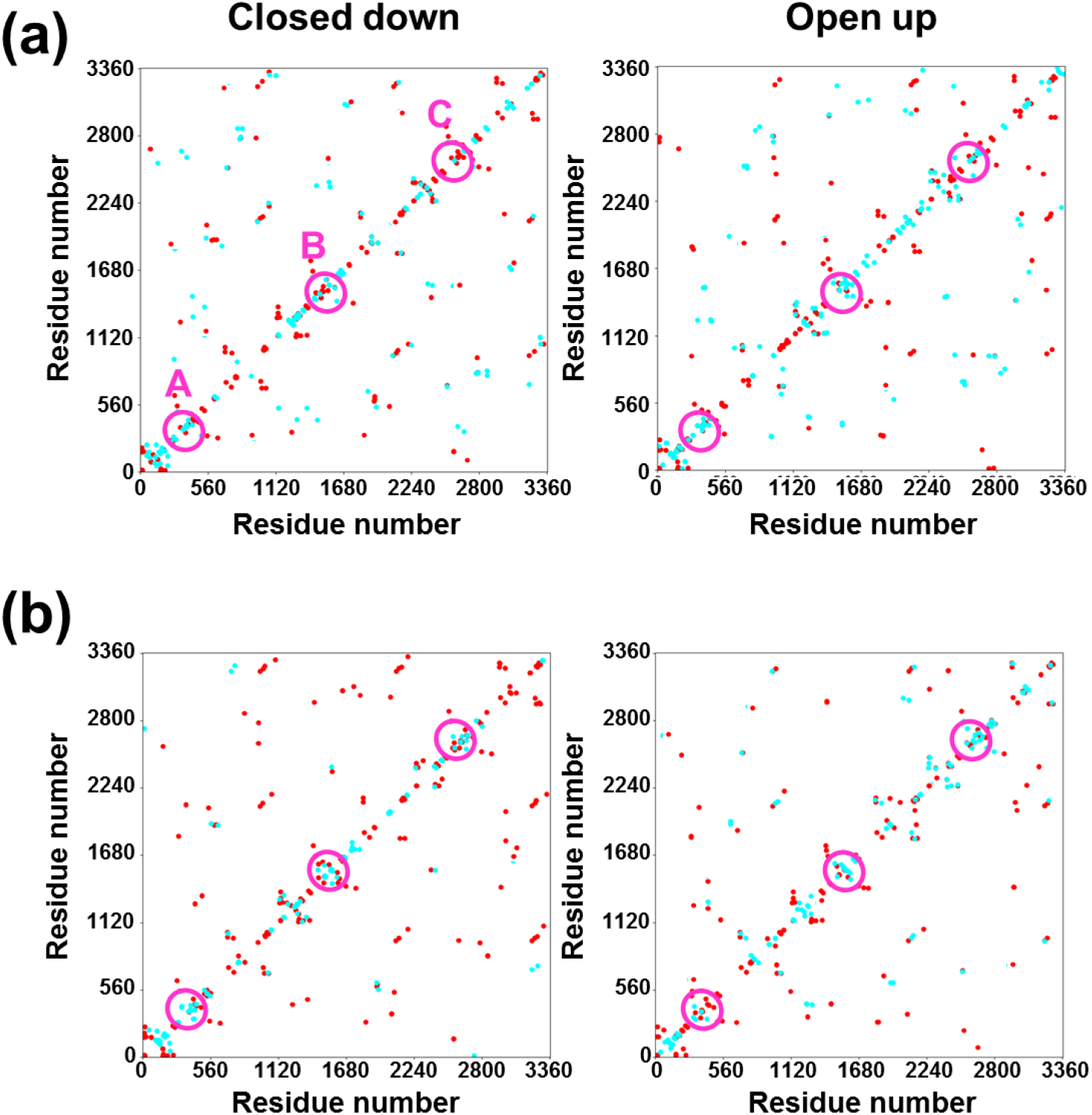
Topology maps for the salt bridges found at the end of the simulations at (a) 373 K and (b) 298 K for the closed down and open up states. The *x*-and *y*-axes represent the residue number containing the positively charged and the negatively charged atoms, respectively. Red dots indicate salt bridges present at the starting structure that are maintained at the end of the simulation (*i.e.* the two residues involved in the salt bridge do not change), while light blue dots correspond to the salt bridges that were not present at the starting conformation (*i.e.* the residue involving the positively or negatively charged atoms change). The empty purple circles indicate the position of the RBD in the three chains of the homotrimer.

The impact of the restrictions on the protein conformation has been further analyzed by considering distortions in the backbone φ,ψ-rotamers. For this purpose, φ,ψ-Ramachandran plots have been depicted for selected snapshots at the beginning (t= 0 ns), middle (t= 75 ns) and end (t= 150 ns) of the simulations. The approximate location and shape of the most favorable low-energy regions in this coordinate system is known to depend on the chemical structure of the residue. In this work, we have applied the classification proposed by Richardson and coworkers (Lovell *et al.*, 2003), according to which φ,ψ-maps of the following residues should be considered separately: 1) glycine (Gly) that is the most flexible residue due to the lack of side chain; 2) proline (Pro) with a singular cyclic structure that prohibits the rotation about the N—C^α^ bond and confines the φ torsion angle at around −60°; 3) residues preceding Pro (pre-Pro), which exhibit a very distinctive φ,ψ-distribution due to the steric restrictions imposed by the neighbor Pro (Moradi *et al.*, 2011); and 4) the general case of the other 18 residues (hereafter, named General) that are largely influenced by the collision among main chain and C^β^ atoms. The energetically allowed φ,ψ-regions for General, Gly, Pro and pre-Pro residues are enclosed by the contours in the maps displayed in Figure S6 (Lovell *et al.*, 2003).

The φ,ψ-maps of General residues for the two states, calculated at 298 and 373 K, are shown in Figure S7a (t= 0 ns) and 7a (t= 75 and 150 ns). It is worth noting that the main part of such residues is located inside favored and allowed regions (blue dots within the contoured zones), while a small part is out but surrounding the allowed regions (red dots over the border or close to the contoured zones). The latter residues, which are found in a similar number for the two states, are modestly strained with a small energetic penalty with respect to the residues inside the favored regions. In addition, a few residues appear at forbidden areas with pronounced steric clashes (red dots over white regions), indicating highly strained conformations. These few disfavored residues appear in the closed state even at the beginning of the simulation (Figure S7a), independently of the temperature, suggesting that they should be associated to the symmetric assembly of the three monomers rather to the heating process. Moreover, highly strained conformations are localized in chains B and C, as is proven in Figure S8 that compares the φ,ψ-maps for the three monomers at the end of the 298 and 373 K trajectories.

This effect is less pronounced for flexible Gly residues, as is shown in the φ,ψ-maps displayed in Figures S7b and 7b. Gly lacks of a side chain, making its φ,ψ values substantially less restricted than those related to other amino acids. Thus, the number of Gly residues localized in highly disfavored conformational regions (red dot over white regions) is very small, regardless of temperature. In this case, the increase in temperature from 298 to 373 K only causes a greater spreading of the blue dots inside the contoured regions, which is associated to the favored and allowed regions. This increment in the flexibility of Gly residue can be associated to structural deformations without significant energy penalties. A similar effect is observed for Pro and, especially, pre-Pro residues, as is shown in Figures S7c-d and S9. Although temperature induces the apparition of a few strained residues for both the closed down and open up states, this straining effect is much less pronounced than for General residues. Besides, the apparition of strained pre-Pro residues is practically undetectable through all trajectories. It is worth noting that, in opposition to Gly, that is very flexible and able to accommodate in many different conformations, the behavior of Pro and pre-Pro has been attributed to the geometric restrictions imposed by the pyrrolidine ring.

**Figure 7.**
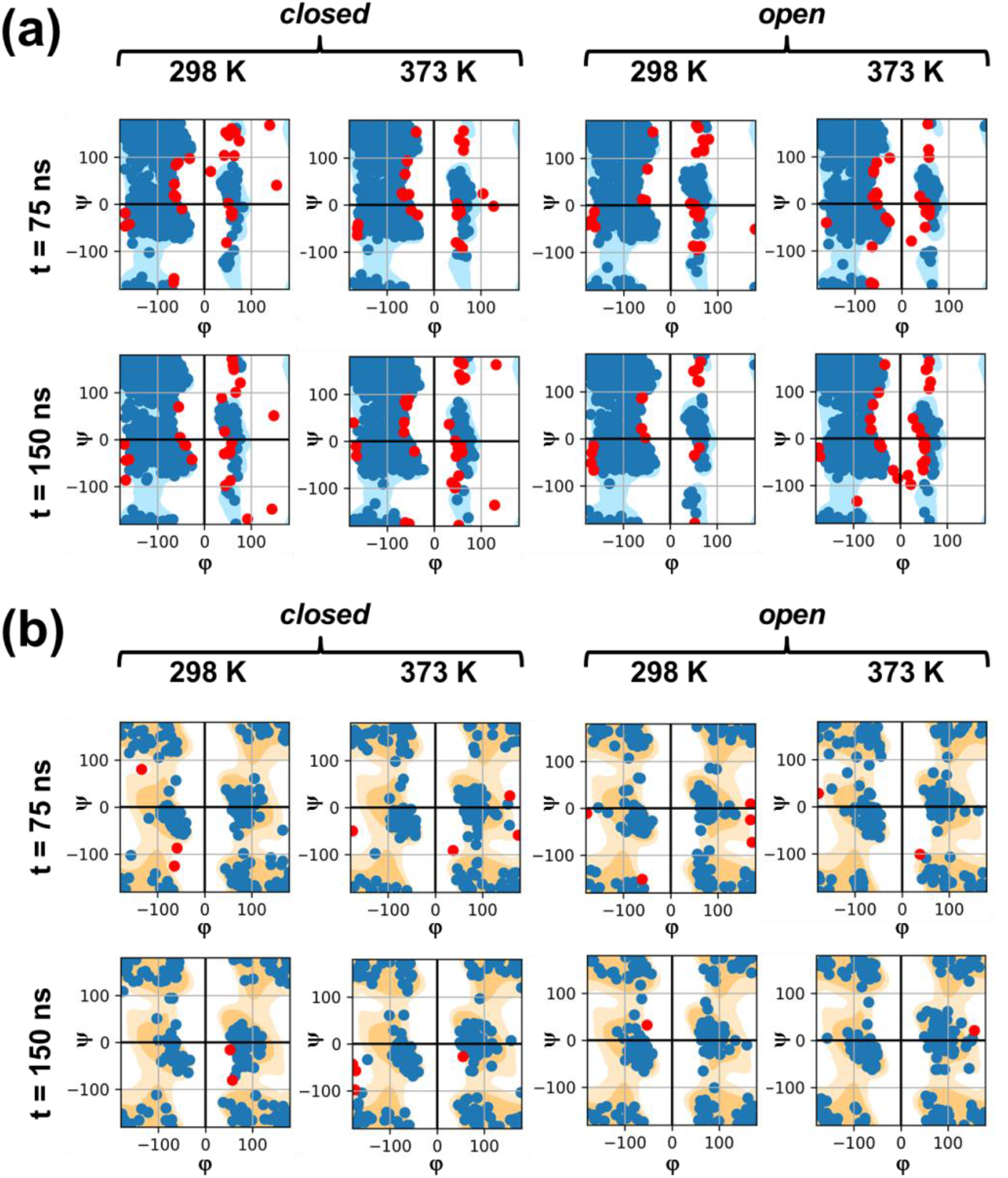
Ramachandran maps showing the φ,ψ angle distributions for (a) General residues (non-Gly, non-Pro and non-prePro) and (b) flexible Gly residues as obtained from snapshots recorded in the middle (t= 75 ns) and at the end (t= 150 ns) of the trajectories conducted at 298 and 373 K for the closed down and open up conformations. Blue dots correspond to residues with allowed conformations (regions enclosed by the contours, see Figure S6), while red dots represent residues with strained (close to regions enclosed by the contours) or very strained (red dots over white regions) conformations.

## Discussion

The stabilization of the closed down conformation with respect to the open up, which is in agreement with experimental observations (Pallesen *et al.* 2017; Walls *et al.*, 2020; Yuan *et al.*, 2017), has been attributed to the attractive inter-monomer interactions since the internal energy of each monomer is similar for the two states. Indeed, the interactions between RBD of a given monomer with the NTDs of neighboring monomers are favored in the closed down state compared to the open up and the interactions in the latter being truncated for the B chain. Moreover, the energy gap between the two states increases with the temperature due to the thermally-induced structural distortions that are more pronounced for the open up state. Instead, water-protein intermolecular interactions favor the open up conformation, which is obviously the consequence of the RBD exposition.

The narrow fluctuations of the A-B, B-C and A-C interchain distances found for the closed down state (Figure 2a) indicate the homogeneous disposition of the three monomers is not perturbed by the temperature, which is consistent with the C3 symmetry imposed for the refinement of the cryo-electron microscopy data (Henderson *et al.*, 2020). In opposition, the interchain distances obtained for the open up state (Figure 2b) reflect not only that the A monomer is closer to C than to B but also that the temperature affects the interchain distances. Thus, the increment of temperature affects the packing of the trimers in the open up state, bringing closer two of the three chains. In spite of this change, the shape of the individual monomers and the whole homotrimer remain practically unaltered with the temperature, as deduced from the calculated R_g_ values (Figure 2c-f).

Analysis of the temporal evolution of the RMSDs at different temperatures (Figure 3a-b) reveals that the thermostability of the two examined conformational states is similar up to 338 K, the close down being more stable than the open up at higher temperatures. The RMSF profiles (Figure 3c-d) reflect that atomic fluctuations associated to thermal instability of the closed down and open up states follow different patterns. More specifically, the higher fluctuations observed for the former state are more localized and less frequent than for the latter state, which exhibits important deviations at larger regions of the protein motifs. This feature is clearly illustrated by ΔRMSF_298-373_ values (Figure 3e-f).

In general, the RMSFs profiles shown in Figure 4 are fully consistent with cryo-electron microscopy observations of SARS-CoV-2 spike, which revealed considerable flexibility and dynamics in the S1 subunit (Kirchdoerfer *et al.*, 2016; Hoffmann, *et al.*, 2020). In addition of the expected flexibility of RBD, which has been extensively investigated in this work by considering the closed down and open up states separately, the NTD flexibility suggested by cryo-electron microscopy structures is confirmed by MD simulations. Another region from S1 that deserves consideration is the one located between the RBD and the S2 subunit, which is usually subdivided in two structurally conserved subdomains identified as SD1 and SD2. These subdomains, especially SD1, act as a hinge point for RBD closed down–to–open up transitions and, therefore, are strongly affected by the breathing movements between the two states, which explain the relatively large fluctuations observed at all studied temperatures.

On the other hand, the preservation of the number of hydrogen bonds with increasing temperature suggests that the glycoprotein spike of SARS-CoV-2 presents some kind of thermal stability, which at a first glance would not justify the inactivation of the virus by temperature. However, further understanding of the latter observation has been achieved by examining the changes in the topology of the hydrogen bonding network induced by the temperature. Thus, a large number of newly formed hydrogen bonds is observed at 373 K. Conversely, native hydrogen bonds were mostly preserved at 298 K for both studied conformational states through the dynamics runs. This difference explains the inactivation of the SARS-CoV-2 when rising the temperature. Although the largest change in the topology is achieved at 373 K, the observation of changes at lower temperatures is probably hindered by the limited length of the trajectories. More specifically, the total inactivation of the virus has been reported at 56 °C after 45 min heating (Jureka *et al.*, 2020), which suggests a very slow kinetics for the conformational and topological changes related to hydrogen bonding reorganization.

Comparison of the topology maps at 298 and 373 K (Figure 5a-b) suggests that, in general, the newly formed hydrogen bonds involve residues that are relatively close to those involved in the native interactions. This feature is consistent with the fact that energy delivered through heating induces changes in the secondary and, even, tertiary structure of the spike glycoprotein but not in its global shape or quaternary structure, as discussed above (Figures 1 and 2). Considering the moderate strength of hydrogen bonds (*i.e.* around –5 kcal/mol), the roughly retention of the shape at 373 K should be related to stronger noncovalent interactions. In fact, the average amount of salt bridges increases ~11% (closed down) / ~18% (open up) when the temperature rises from 298 to 373 K, supporting that salt bridges are responsible for the global shape retention observed at 373 K.

Overall, analyses of the temperature influence on both hydrogen bonds (weak directional interactions) and salt bridges (strong non-directional interactions) allow us to propose a molecular mechanism for the thermal inactivation of the virus. Such mechanism consists on differentiating the biological stability from the structural stability. The preservation of a large number of salt bridges and the formation of new ones upon increasing temperature are responsible of the structural stability in terms of protein global shape. Conversely, the deterioration of the hydrogen bonding network appears as the main reason for the functional inactivation of the virus upon heating. This mechanism is summarized in Scheme 1, which depicts the drastic conformational changes experienced by the NTD and RBD of chain B when temperature increases from 298 K to 373 K, whereas the overall shape of the spike is roughly preserved.

**Scheme 1.**
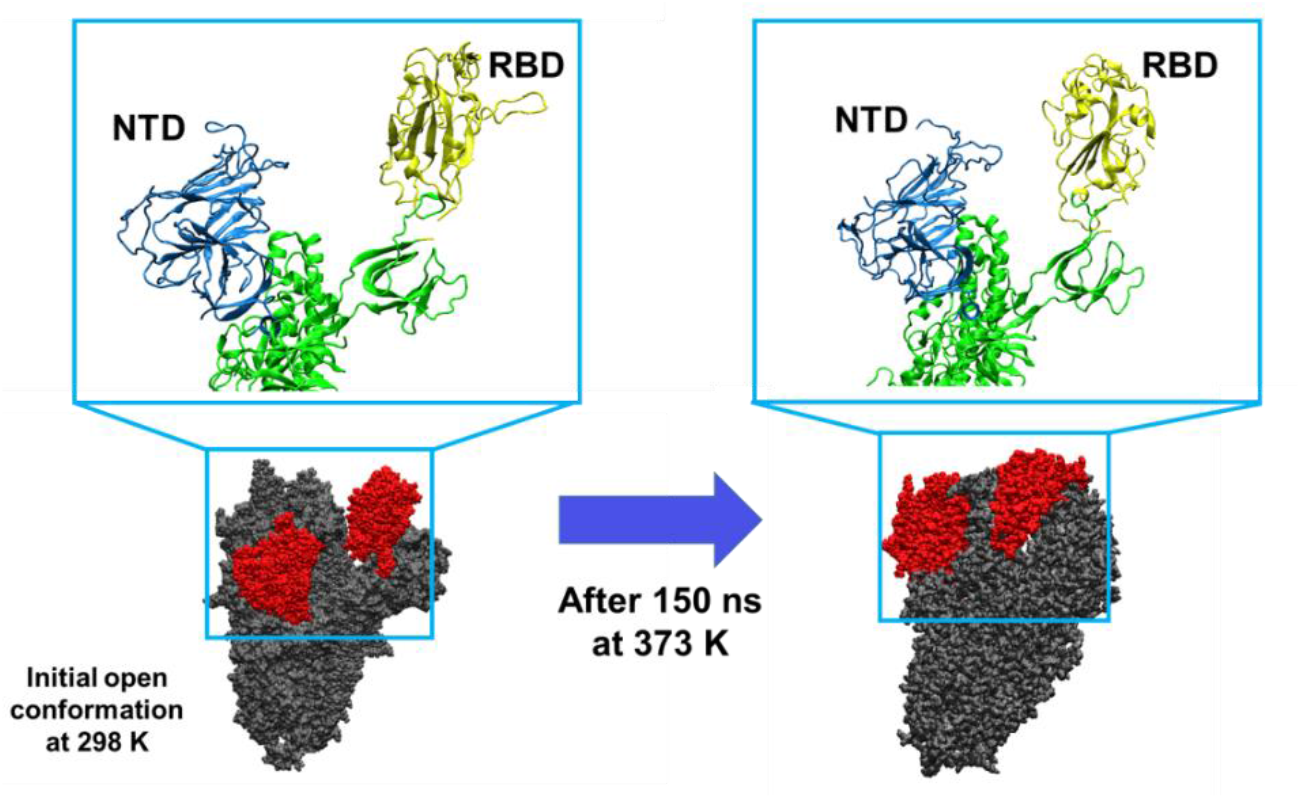
The increase of temperature produces drastic conformational changes in the S1 subunit of the heterotrimeric spike protein (explicitly sketched for chain B), affecting also the orientation of the regions containing the NTD and RBD domains (in red). These changes are proposed to inactivate the virus by modifying the binding area to the cellular receptor, even though the general shape of the whole proteins remains roughly unaltered (grey).

The loss of most of the initial hydrogen bond interactions due to the increase in temperature facilitates new interactions with the surrounding active groups. However, the salt bridges structure, still preserved, restricts the degrees of freedom for establishing new interactions. In general, the geometric constrains imposed by salt bridges limit the formation of new hydrogen bonds among residues located relatively close to those that make functional the infective form. The strength of salt bridges restricts the flexibility of the protein domains, limiting the molecular fluctuations and, consequently, the range of residues that can be involved in suitable directional interactions, involving only those that are spatially close. Such specific interactions result in the new hydrogen bond network that defines the topology of the inactive form of the virus.

The φ,ψ-maps displayed in Figures 7 and S7–S9 are fully consistent with the mechanism proposed in Scheme 1. Thus, the General residues, which are able to form hydrogen bonds and salt bridges, experience not only the largest conformational changes but also, in some cases, adopt strained conformations, as suggested by the topology maps calculated for those interactions. Such findings, open new avenues to develop strategies to inactivate the virus, by targeting specific areas of the homotrimer in order to misbalance, remodel or cleave the hydrogen bonds that make the virus functional (*i.e.* by using chaotropic agents or surfactants that make feasible the disruption of hydrogen and salt bridges). Furthermore, it allows the development of active devices able to deliver energy that can cleave and reorganize hydrogen bonds, for instance, through ultrasounds, irradiation or by using nanoparticles as nanosources of heat able to induce a local increment of temperature by using the local surface resonant plasmon effect, a phenomenon that occurs when light interacts with matter at specific wavelengths. The effectiveness of such local thermal treatment could be increased by controlling the intensity of the light source, the irradiation time lapse and the functionalization of nanoparticle designed to maximize the interaction with the virus areas where the higher number of hydrogen bonds is exposed.

In summary, our results suggest that temperature induces conformational changes on the S1 subunit of the spike glycoprotein of SARS-CoV-2 that remodel the internal hydrogen bonding structure. However, the impact of temperature, even at 373 K, is not strong enough to induce a significant modification in the global shape of the spike. Those conformational changes, are much more pronounced in the state where the binding domain is accessible and the virus is infective (open up state) than when the binding domain is retracted (closed down state). This effect has been associated to a drastic modification in the hydrogen bonding topology, which particularly affects the recognition functionality of the receptor binding domain. Such network reorganization is triggered by the energy delivered through heat that allows the massive cleavage and remodeling of a significant amount of hydrogens bonds. Conversely, the salt bridges topology remains much less altered, allowing to maintain the main structure that defines the shape of the spike. The deep knowledge about such inactivation mechanism facilitates the development of new strategies intended to inactivate the virus through the destruction or modification of such hydrogen bonds by chaotropic agents and surfactants or through physical treatments able to selectively target such labile hydrogen bond structures.

## Methods

### Construction of the molecular models

Cryoelectron microscopy structures of the homotrimeric spike glycoprotein of SARS-CoV-2 in the closed down (PDBid: 6vxx) and open up (PDBid: 6vyb) conformational states, solved at 2.80 Å and 3.20 Å, respectively (Walls *et al.*, 2020), were taken from the Protein Data Bank and used to prepare the initial conformations. The missing residues (*i.e.* 44-55, 88, 89, 118-139, 147-159, 217-236, 417-429, 445-462, 476, 595-614, 651-653, 802-828, 1162-1175, 1236-1258, 1268-1280, 1292-1294, 1307-309, 1338-1357, 1550-1556, 1562-1585, 1610-1616, 1716-1735, 1772-1783, 1923-1948, 2283-2296, 360-2380, 2389-2401, 2459-2479, 2661-2263, 2671-2677, 2687-2706, 2837-2856, 2893-2905 and 3044-3071) were incorporated using the Modeler algorithm (Webb and Sali, 2016) implemented in the UCSF Chimera program (Pettersen *et al.*, 2004) and the Z-DOPE (Discrete Optimized Protein Energy), statistical potential based for the choice of best model (*i.e.* that with the lowest Z-DOPE) for each conformational state, among the generated ones.

The homotrimeric protein models were then submerged in a previously equilibrated water box of 166.04 × 175.56 × 200.82 nm^3^ for the closed down conformational state and of 161.05 × 183.59 × 213.39 nm^3^ for the open up one. Any water molecule that overlapped with any of the atoms belonging to the protein spike model was removed and, finally, a total of 154517 and 168125 water molecules were kept for the closed down and open up state, respectively. The models were completed by inserting randomly Na^+^ and Cl^−^ ions until reaching a 0.15 M NaCl concentration to reproduce physiological conditions. Then, the two models were processes with the LEaP program (Case *et al.*, 2005) to add hydrogen atoms to the protein and to generate Amber topology files and coordinates files. Accordingly, the models used to represent the closed down and the open up states of the homotrimeric spike protein presented 516706 and 558601 explicit atoms, respectively.

### Computational details

All simulations were performed using the AMBER 18 simulation suite (Case *et al.*, 2018). Protein atoms were modeled using the Amber ff14SB force field (Maier *et al.*, 2015), the glycan atoms included in the cryo-EM coordinates were modeled using the Glycam06 force field (Kirschner *et al.*, 2008), and water atoms were modeled using the TIP3P force field (Jorgensen *et al.*, 1983).

Equilibration calculations were started by relaxing the protein regions filled with the UCSF Chimera program (Pettersen *et al.*, 2004), which was achieved by applying the Limited-memory BroydenFletcher-Goldfarb-Shanno quasi-Newton algorithm methodology to the new added residues meanwhile the rest of the system was kept frozen. Next, the whole system was submitted to 2500 steps of full conjugate gradient minimization to relax conformational and structural tensions.

The Langevin dynamics method (Izaguirre *et al.*, 2001) was used to heat the system and to rapidly equilibrate its pressure and temperature. The relaxation times used for the coupling were 5 ps for both temperature and pressure. The temperature was increased from 0 K to 298 K using 60 ps simulation in the NVT ensemble, using an integration time step of 1 fs and keeping the pressure at 1.034 bar. Then, 1 ns in the NPT ensemble were conducted at 298 K to relax the structure and the density (integration step: 2 fs; pressure:1.034 bar). The last snapshot of this relaxation was used not only as starting point of the NPT production trajectory at 298 K but also as starting point of 0.25 ns NPT-MD simulation to bring the temperature to 310 K (integration step: 2 fs; pressure:1.034 bar). The last snapshot was used as starting point for the NPT production trajectory at such temperature and as starting point for the NPT-MD used to bring the temperature to 324 K. The same process was used to generate starting points for the production simulations 338, 358 and 373 K.

Production NVT-MD trajectories at 298, 310, 324, 338, 358 and 373 K were 150 ns each. Replica simulations were performed at all the indicated temperatures by changing the initial velocities. Accordingly, a total of 3.6 μs (150 ns × 2 conformational states × 6 temperatures × 2 replicas) were simulated for the studied system.

Non-bonding pairs list was updated every 10 steps. Periodic boundary conditions were applied using the nearest image convention and the atom pair cut-off distance used to compute the van der Waals interactions was set at 10.0 Å. In order to avoid discontinuities in the potential energy function, non-bonding energy terms were forced to slowly converge to zero, by applying a smoothing factor from a distance of 12.0 Å. Beyond cut-off distance, electrostatic interactions were calculated by using Particle Mesh of Ewald (Toukmaji *et al.*, 2000).

## Acknowledgements

Authors acknowledge PRACE for awarding us access to Joliot-Curie at GENCI@CEA(Irene), France, through the “PRACE support to mitigate impact of COVID-19 pandemic” call, Agència de Gestió d’Ajuts Universitaris i de Recerca (2017SGR359 and 2017SGR373) and B. Braun Surgical, S.A.U. for financial support. Support for the research of C.A. was received through the prize “ICREA Academia” for excellence in research funded by the Generalitat de Catalunya.

## Competing interests

There is no competing interest.

## Supplementary Figures

**Figure S1.**
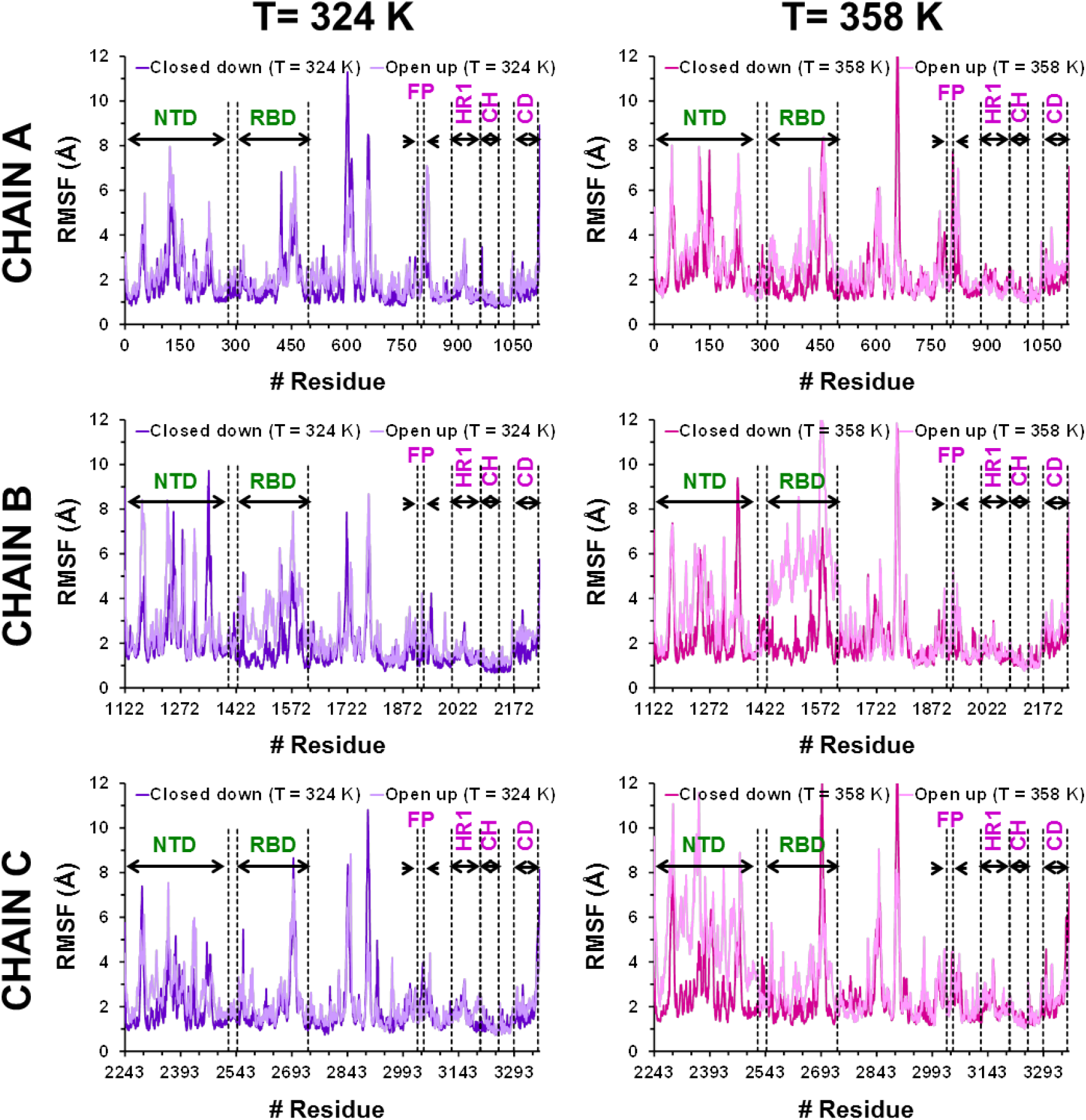
For the three chains of the homotrimeric spike protein in the closed down and the open up states: root mean square fluctuation (RMSF) analyses calculated using all atoms for the molecular dynamics simulations at 324 K (left) and 358 K (right). The following domains are indicated in each graphic: the N-terminal and the receptor binding domain (NTD and RBD, respectively) from S1 subunit, and the fusion peptide (FD), the heptapeptide repeat sequence 1 (HR1), the central helix (CH), the connector domain (CD) from S2 subunit.

**Figure S2.**
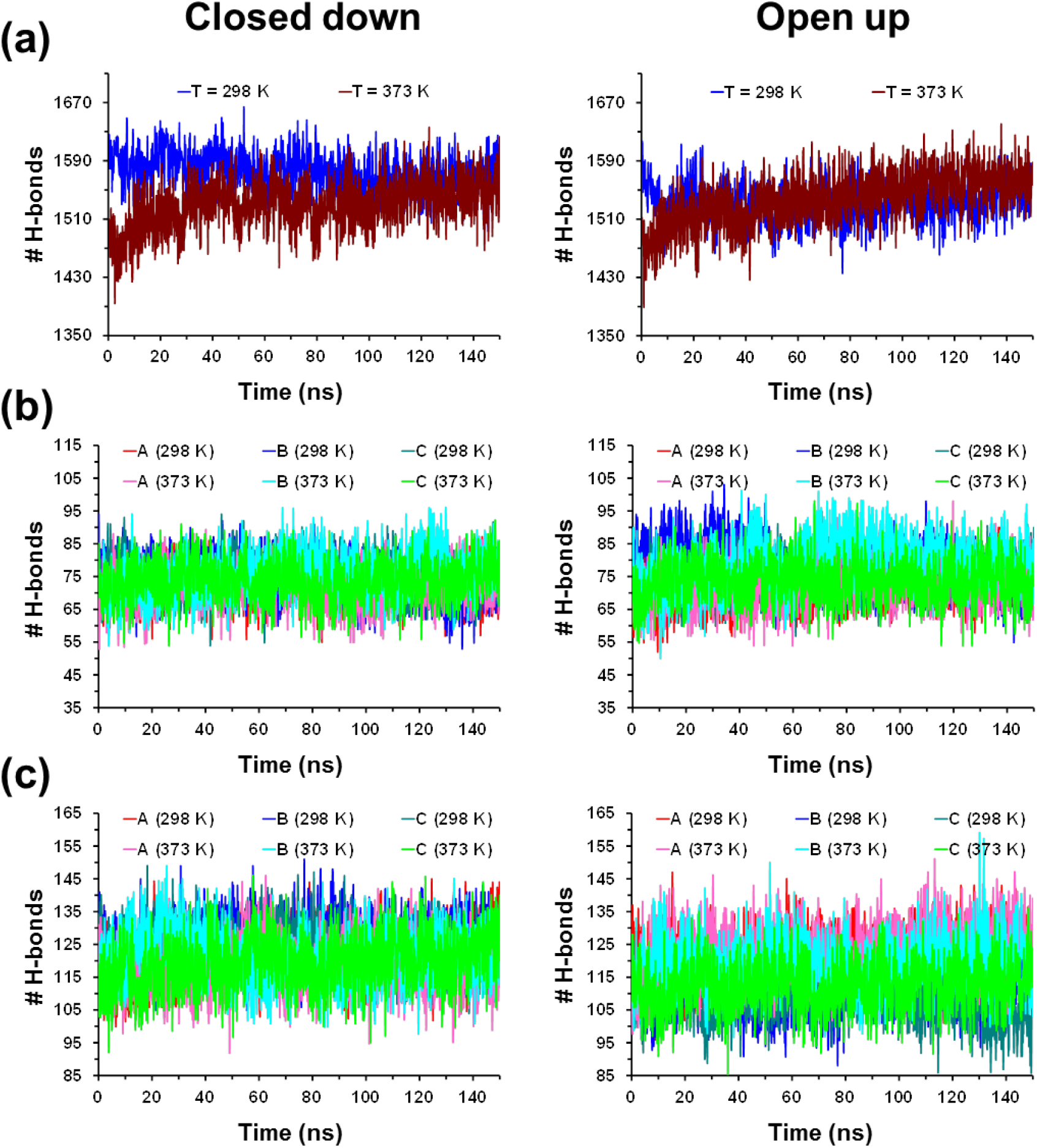
For the homotrimeric spike protein in the closed down (left) and open up (right) states: (a) temporal evolution of the total number of hydrogen bonds at 298 and 373 K; (b) temporal evolution of the number of hydrogen bonds involving residues from the RBD of chain A, B and C at 298 and 373 K; and (c) temporal evolution of the number of hydrogen bonds involving residues from the NTD of chain A, B and C at 298 and 373 K. The geometric parameters and the cut-off values used to define a D–H···A hydrogen bond (where A is an acceptor atom, D a donor heavy atom and H a hydrogen atom) are the ∠DHA angle, which must be greater than 135°, and the D···A distance, that must be less than 3.0 Å.

**Figure S3.**
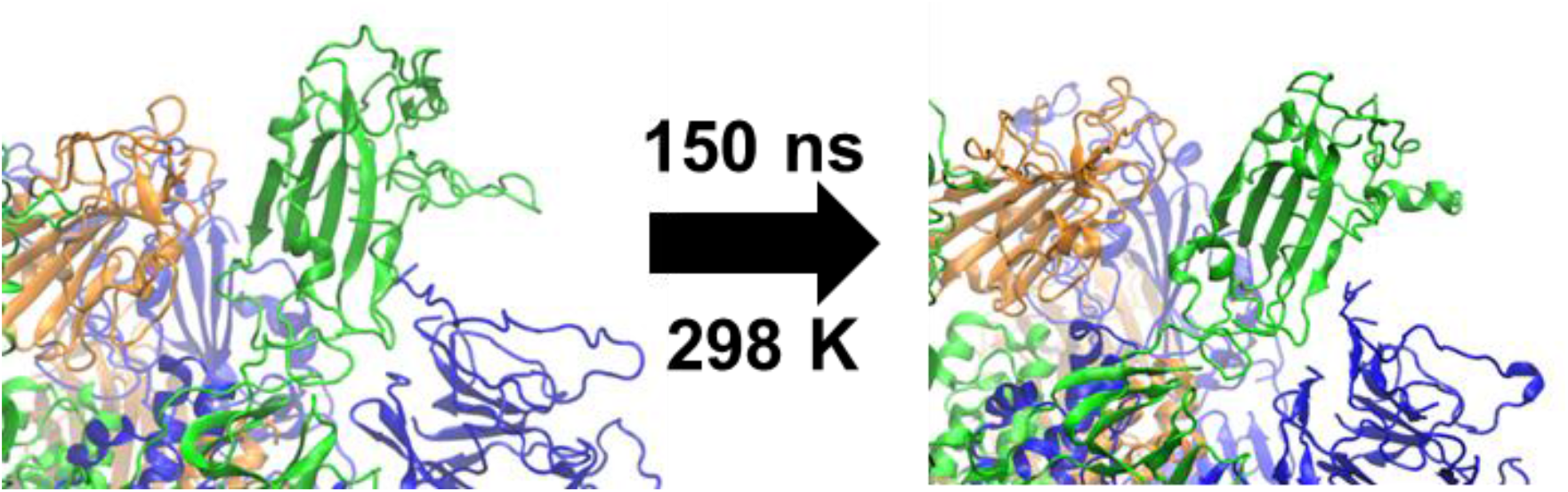
Representation of the region containing the RBD domains at the beginning and the end of the simulation for the open up state at 298 K.

**Figure S4.**
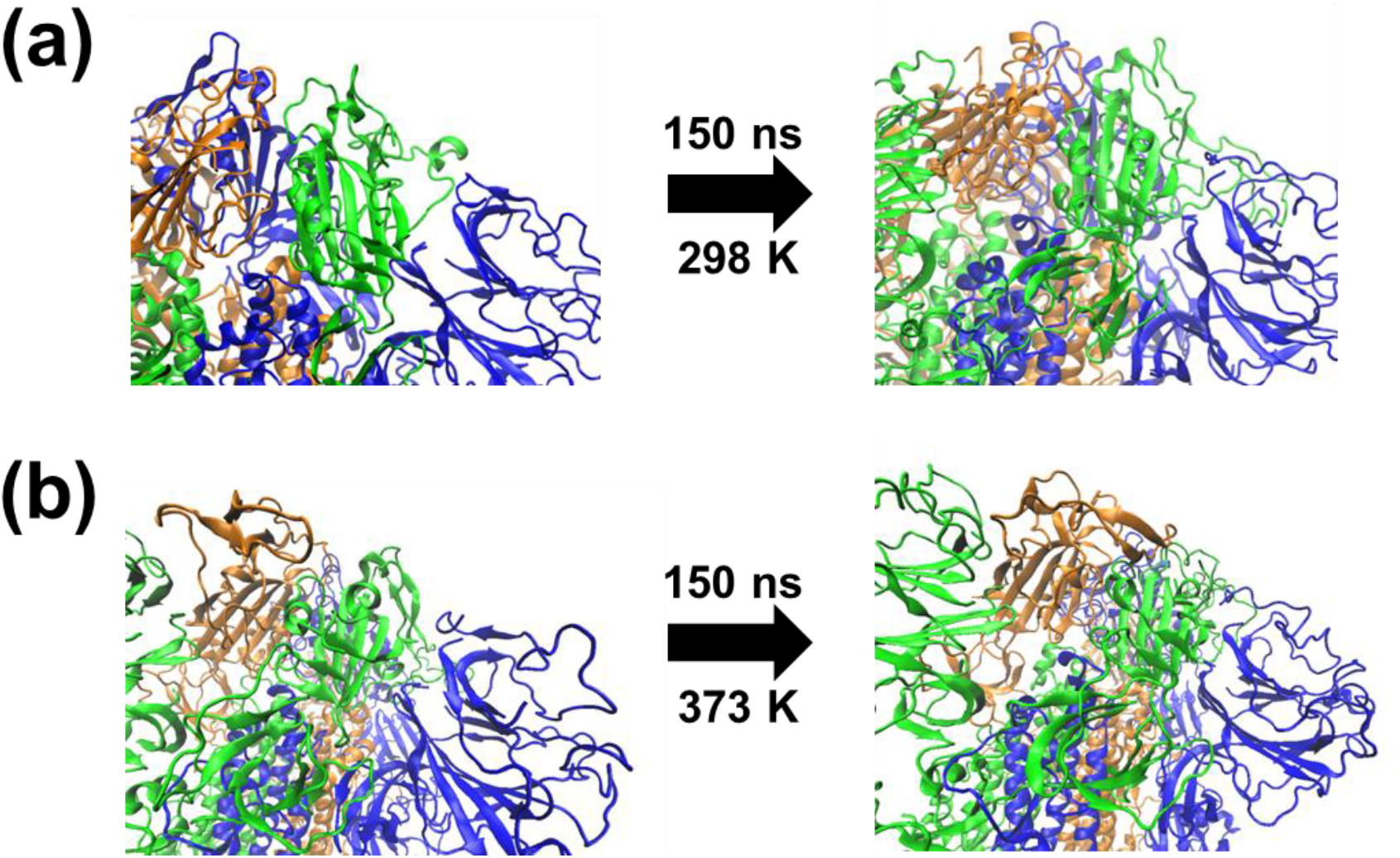
Representation of the region containing the RBD domains at the beginning and the end of the simulation for the closed down state at (a) 298 K and (b) 373 K.

**Figure S5.**
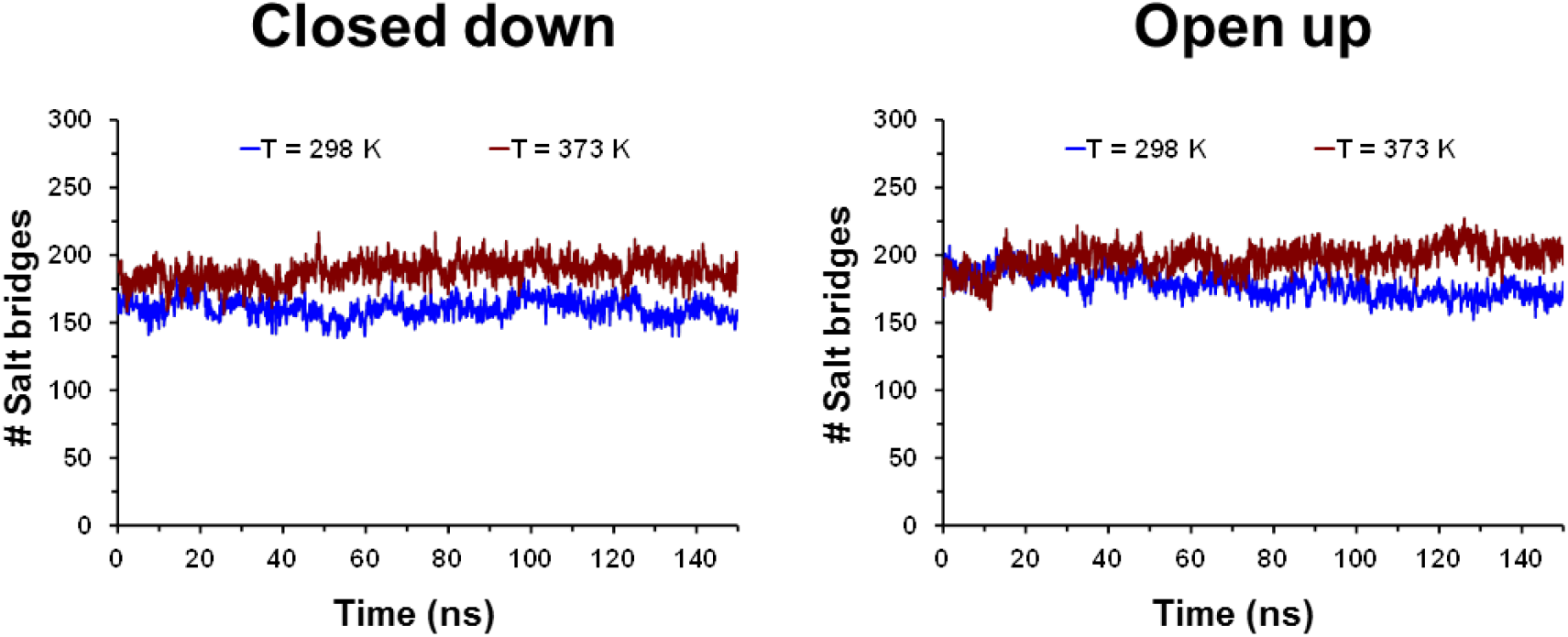
Temporal evolution of the total number of salt bridges at 298 and 373 K for the homotrimeric spike protein in the closed down (left) and open up (right) states.

**Figure S6.**
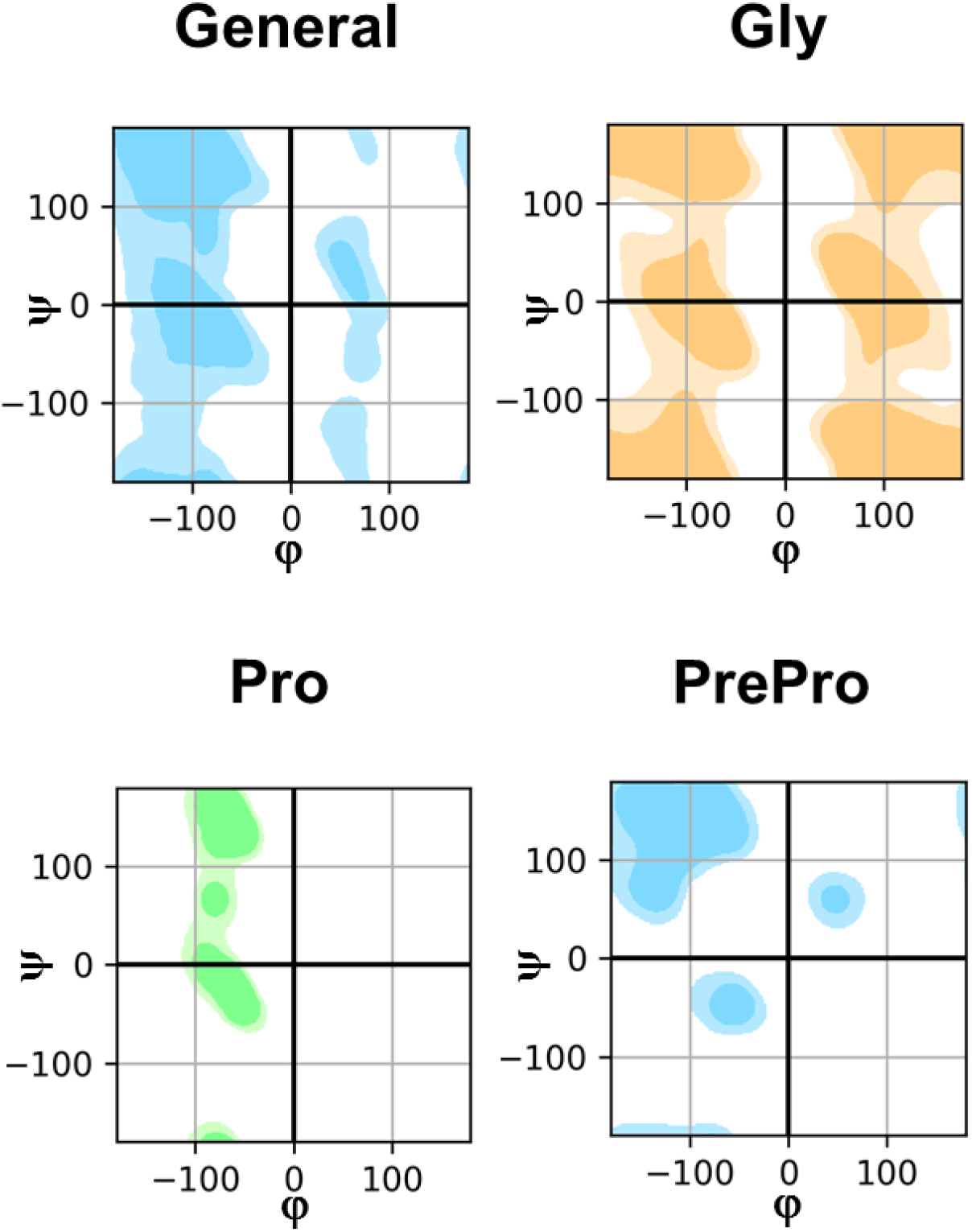
φ,ψ-Ramachandran maps indicating the allowed regions (enclosed by the contours) for General, Gly, Pro and PrePro residues.

**Figure S7.**
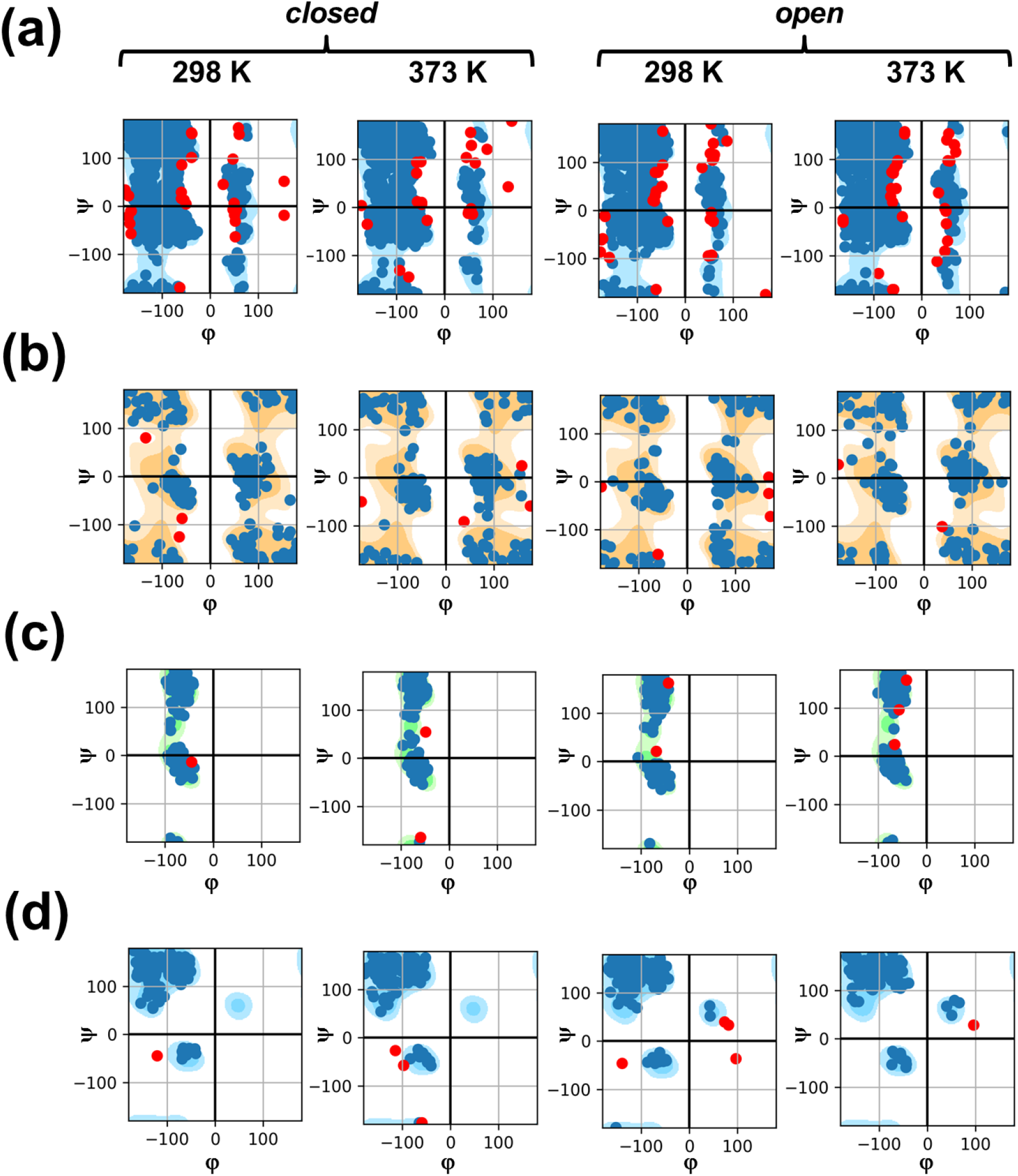
Ramachandran maps showing the φ,ψ angle distributions for (a) General (non-Gly, non-Pro and non-PrePro), (b) Gly, (c) Pro and (d) PrePro residues at the beginning of the trajectories (t= 0 ns) conducted at 298 and 373 K for the closed down and open up conformations. Blue dots correspond to residues with allowed conformations (regions enclosed by the contours, see Figure S6), while red dots represent residues with strained (close to regions enclosed by the contours) or very strained (red dots over white regions) conformations.

**Figure S7.**
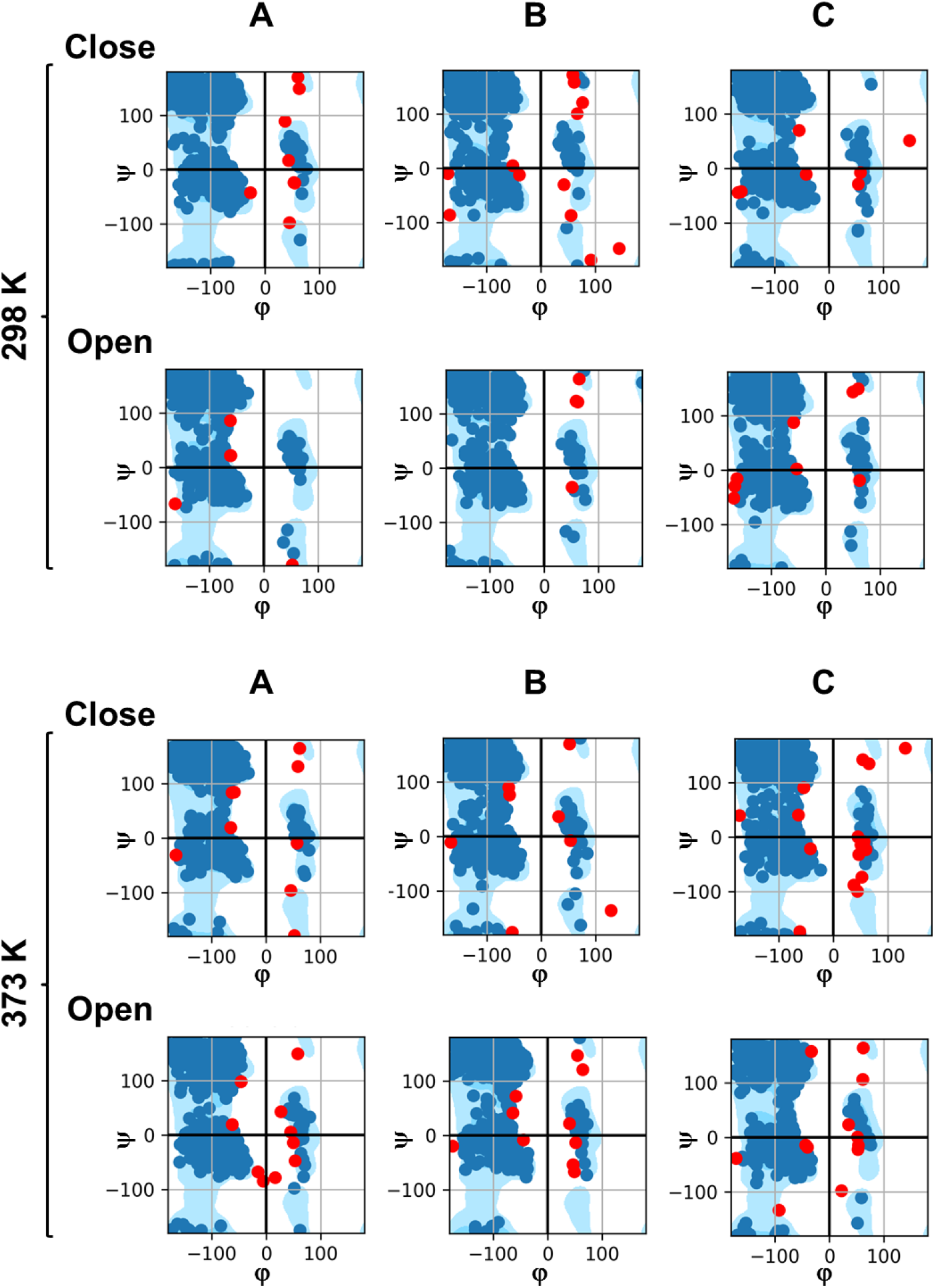
Ramachandran maps showing the φ,ψ angle distributions for General residues (non-Gly, non-Pro and non-PrePro) obtained from snapshots recorded at the end of the trajectories conducted at 298 and 373 K for the closed down and open up conformations. Distributions are depicted considering monomers A, B and C separately. Blue dots correspond to residues with allowed conformations (regions enclosed by the contours, see Figure S6), while red dots represent residues with strained (close to regions enclosed by the contours) or very strained (red dots over white regions) conformations.

**Figure S8.**
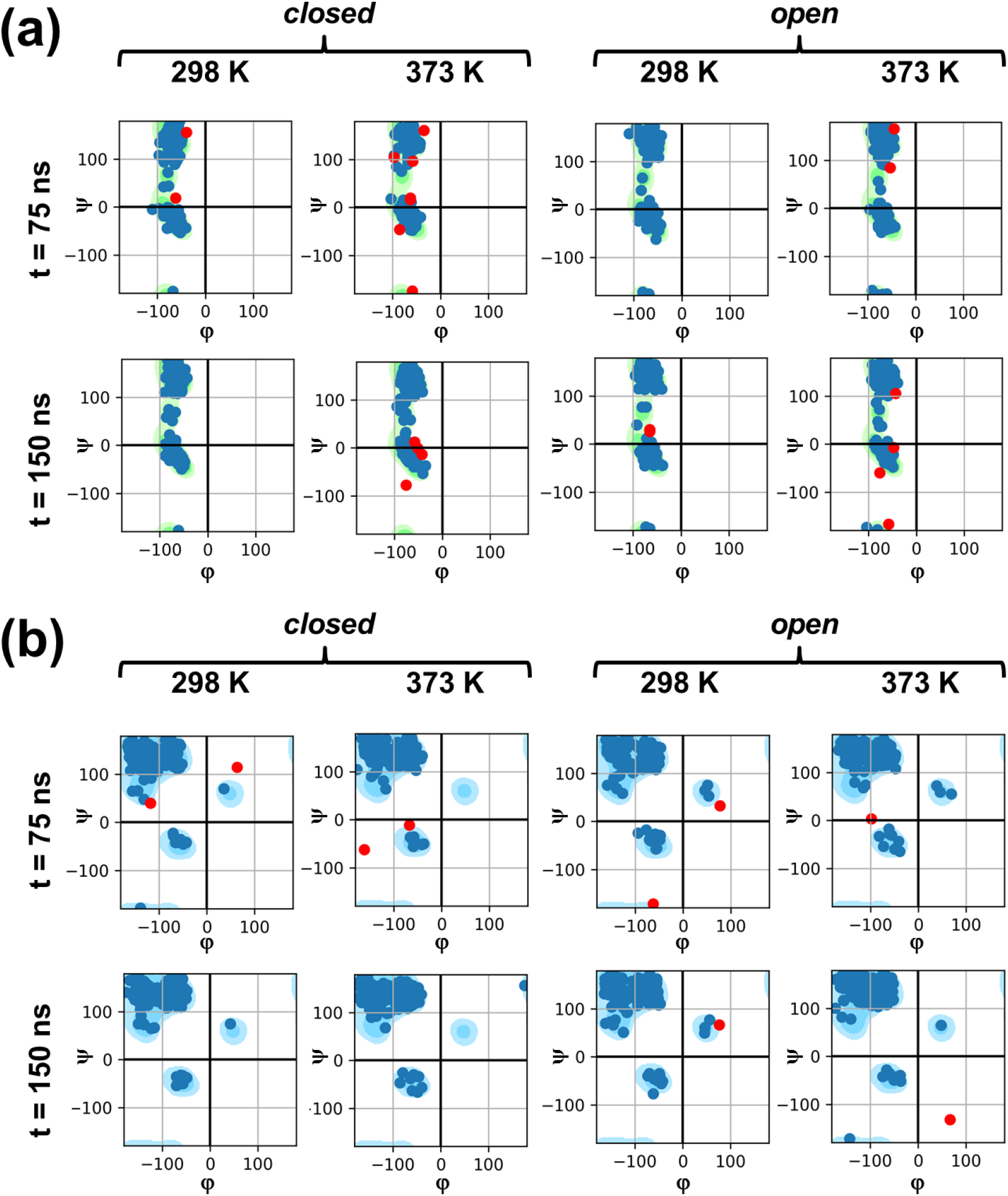
Ramachandran maps showing the φ,ψ angle distributions for restrained (a) Pro and (b) PrePro residues as obtained from snapshots recorded in the middle (t= 75 ns) and at the end (t= 150 ns) of the trajectories conducted at 298 and 373 K for the closed down and open up conformations. Blue dots correspond to residues with allowed conformations (regions enclosed by the contours, see Figure S6), while red dots represent residues with strained (close to regions enclosed by the contours) or very strained (red dots over white regions) conformations.

## Notes

### Competing Interest Statement

The authors have declared no competing interest.

